# WHOLE-GENOME SEQUENCING AND *DE NOVO* ASSEMBLY OF A 2019 NOVEL CORONAVIRUS (SARS-COV-2) STRAIN ISOLATED IN VIETNAM

**DOI:** 10.1101/2020.06.12.149377

**Authors:** Le Tung Lam, Nguyen Trung Hieu, Nguyen Hong Trang, Ho Thi Thuong, Tran Huyen Linh, Luu Thuy Tien, Nguyen Thi Ngoc Thao, Huynh Thi Kim Loan, Pham Duy Quang, Luong Chan Quang, Cao Minh Thang, Nguyen Vu Thuong, Hoang Ha, Chu Hoang Ha, Phan Trong Lan, Truong Nam Hai

## Abstract

The pandemic COVID-19 caused by the zoonotic virus SARS-CoV-2 has devastated countries worldwide, infecting more than 4.5 million people and leading to more than 300,000 deaths. Whole genome sequencing (WGS) is an effective tool to monitor emerging strains and provide information for intervention, thus help to inform outbreak control decisions. Here, we reported the first effort to sequence and *de novo* assemble the whole genome of SARS-CoV-2 using PacBio’s SMRT sequencing technology in Vietnam. We also presented the annotation results and a brief analysis of the variants found in our SARS-CoV-2 strain, which was isolated from a Vietnamese patient. The sequencing was successfully completed and *de novo* assembled in less than 30 hours, resulting in one contig with no gap and a length of 29,766 bp. All detected variants as compared to the NCBI reference were highly accurate as confirmed by Sanger sequencing. The results have shown the potential of long read sequencing to provide high quality WGS data to support public health responses, and advance understanding of this and future pandemics.

## INTRODUCTION

COVID-19, a disease caused by SARS-CoV-2 (Zhou *et al.*, 2020), first emerged in Hubei province, China, in December 2019 and was declared a global pandemic by WHO in March 2020 (WHO, 2020). As of May 2020, more than 4.5 million cases of COVID-19 have been recorded worldwide, resulting in more than 300,000 deaths (John Hopkins University, 2020; Ministry of Health, 2020). COVID-19 is affecting 213 countries and territories around the world, forcing unprecedented measures from every government including border closure, halting of trade, lockdown of cities and social distancing. (WHO, 2020; Ministry of Health, 2020). In Vietnam, there are currently 320 cases that are reported to be positive with SARS-CoV-2; and as of 18^th^ May, there are more than 70,000 people in involuntary isolation (Ministry of Health, 2020).

SARS-CoV-2 is an enveloped virus that carries an approximately 30 kilobases (kb) singlestranded positive-sense RNA genome, and belongs to the genus *Betacoronavirus* in the family *Coronaviridae* (Coronaviridae Study Group of ICTV, 2020) in the same species as SARS-CoV and SARS-related bat CoVs (Lu *et al.*, 2020; Chan *et al.*, 2020). Patterns of spread indicated that SARS-CoV-2 can be transmitted person-to-person, and may be more transmissible than SARS-CoV (Li *et al.*, 2020; Chen *et al.*, 2020). The SARS-CoV-2 genome was reported to possess 14 open reading frames (ORF) encoding 27 proteins (Wu *et al.*, 2020). The release of the nucleotide sequences of SAR-CoV-2 just two weeks after the discovery of the first reported case contributed to the rapid establishment of possible origins and in-depth studies of this virus (Lu *et al.*, 2020; Ceraolo *et al.*, 2020). As of May 2020, more than 30,000 genome sequences of SAR-CoV-2 from many countries have been sequenced and uploaded to international, open access databases including the GISAID Initiative (Elbe & Buckland-Merrett, 2017) and GenBank (Benson *et al.*, 2015).

Whole genome sequencing (WGS) is a robust tool to provide a detailed picture of the sources of infection and transmission chains, and thus help to inform outbreak control decisions (Gwinn *et al.*, 2019). Recently, genomic information has been regularly utilized to reconstruct several outbreaks of zoonotic viral pathogens (Oude *et al.*, 2020; Quick *et al.*, 2016). Sharing and combining genome sequence data during viral outbreaks is now recommended as an integral part of outbreak response. However, among the genome sequences submitted to GISAID, many assemblies are low quality. They can be fragmented or have multiple bases gaps that break ORF, preventing correct analysis of identification and evolution of the virus (Ashby, 2020). These are mainly caused by the short amplicon method’s limitation of the second-generation sequencing (SGS) technologies, most notably Illumina which is being preferred by many laboratories worldwide (Goodwin *et al.*, 2016; Shendure *et al.*, 2017, Ashby, 2020). Meanwhile, singlemolecule, real-time (SMRT) sequencing developed by Pacific Biosciences (PacBio) offers significantly long read lengths with high accuracy, overcoming errors caused by problematic genomic regions (Rhoads & Au, 2015). Most importantly, the highly-contiguous *de novo* assemblies using PacBio sequencing can close gaps in current reference assemblies and characterize structural variation in genomes (Rhoads & Au, 2015). The robustness of this method has been increasingly applied for whole genome SAR-CoV-2 sequencing (Gonzalez-Reiche *et al.*, 2020; CDC, 2020).

As a contribution to the global efforts to track and trace the ongoing COVID-19 pandemic, here we report a method for whole genome sequencing of SARS-CoV-2 using PacBio’s SMRT sequencing technology and the *de novo* assembly and annotation results. We also present a brief analysis of the variants found in our SARS-CoV-2 strain, which was isolated from a Vietnamese patient in Ho Chi Minh City.

## MATERIALS AND METHODS

### SARS-CoV-2 patient information

Samples from a patient denoted as A was enrolled in this study. The patient is a 20-year-old woman returning to Vietnam on March 14^th^, 2020 from the United States (Pensylvania – Philadelphia), transited in Taiwan and arrived in Ho Chi Minh City on flight number BR 395 two days later. The patient was asymptomatic, and was quarantined at the airport. She did not have any symptoms upon arrival but the oropharyngeal (OP) and nasopharyngeal (NP) swabs were still collected as required. SARS-CoV-2 virus infection was detected by real-time reverse transcription PCR (RT-PCR) at the Pasteur Institute in Ho Chi Minh City.

### Virus cultivation

The Vero E6 cell lines were investigated for their susceptibility to SARS-CoV-2 and used for virus isolation in this study. They were cultured in Dulbecco modified Eagle medium (DMEM) supplemented with heat inactivated fetal bovine serum (10%) and antibiotic and grown in a humidified 37°C incubator with 5% CO_2_. Confluent cells were maintained at 37°C in 25 cm^2^ flask containing 5 mL maintenance medium supplemented with 16 mg/mL trypsin-TPCK, 100 U/mL penicillin, and 100 μg/mL streptomycin. They were infected with filtered specimens. After infection, all flasks were observed for cytopathic effects (CPE). Once the CPE was occurred, cell culture supernatants were collected for virus detection and quantification by rRT-PCR, harvested and divided into some aliquot tubes. If no CPE was observed after 72 hours, cell lines negative for indicators of viral replication were blind-passages twice.

### RNA extraction and cDNA synthesis

A 560 μL volume of lysis buffer containing guanidinium and carrier-RNA (QIAGEN, Hilden, Germany) was added to 200 μL supernatant from cell cultures. The sample was removed from the BSL-3 to a BSL-2 laboratory, where it underwent nucleic acid extraction using the QIAamp Viral RNA mini kit (QIAGEN, Hilden, Germany) according to the manufacturer’s instruction. A 50 μL volume of elute was used for removing contaminating DNA from RNA preparation by Turbo DNA (Invitrogen, Carlsbad, CA, USA). A 8 μL volume of RNA, 1 μL of 50 ng/μL random hexamers (Invitrogen, Carlsbad, CA, USA), and 1 μL of 10 μM de-oxy-nucleoside triphosphates (Invitrogen, Carlsbad, CA, USA) were treated for 5 min at 65°C and added to 10 μL reverse transcription master mix containing 200 U/μL M-MLV RT enzyme (Invitrogen, Carlsbad, CA, USA). After incubation at 25°C for 10 min, 50°C for 50 min, then 85°C for 15 min, a 1 μL Rnase H was added and incubated at 37°C for 20 min. NEBNext Ultra II Non-directional RNA Second strand synthesis module (New England BioLabs, Ipswich, MA, USA) enzyme mix was used for generating double strand DNA from first strand cDNA after incubation at 16°C for 1 hour.

### Library preparation

The dsDNA products were purified by Agencourt AMPure XP beads (Beckman Coulter, Brea, CA, USA). Before library preparation, input cDNA was quantified using the Qubit fluorometer and Qubit High Sensitivity (HS) DNA assay reagents (ThermoFisher Scientific, Waltham, MA, USA). The input cDNA length was accessed using the Agilent 2100 Bioanalyzer system (Agilent, Santa Clara, CA, USA). SMRTbell Libraries was prepared using Express Template Prep Kit 2.0 with low DNA input (50 ng) (Pacific Biosciences, Menlo Park, CA, USA) for sequencing on the PacBio SEQUEL system according to the manufacturer’s instruction. Briefly, DNA damage was repaired using the SMRTbell Damage Repair Kit - SPv3 (Pacific Biosciences, Menlo Park, CA, USA), the product was introduced for End-repair/A-tailing followed by ligation to PacBio’s adapter. The library was purified one time with 1.2 volumes of AMPure PB beads (Beckman Coulter, Brea, CA, USA). The size and amount of library was checked again using the Bioanalyzer 2100 system and the Qubit fluorometer with Qubit High Sensitivity (HS) DNA assay reagents, respectively. The library was then applied on a SMRT Cell (Pacific Biosciences, Menlo Park, CA, USA). Total time for library preparation was 8 hours.

### Whole-genome sequencing of a SARS-CoV-2 strain with PacBio SMRT

The library was bound to polymerase using Sequel I Binding Kit 3.0, (Pacific Biosciences, Menlo Park, CA, USA) and purified by Ampure PB beads. DNA Control Complex 3.0 and Internal Control Kit 3.0 (Pacific Biosciences, Menlo Park, CA, USA) were used to control the sequencing procedure. The run design was created by Sample Setup software included in the SMRTLink portal v5.1 with insert size of 1200 basepairs (bp), immobilization time of 2 hours, pre-extension time of 2 hours and movie time of 10 hours.

### *De novo* assembly

The sequencing signals were processed, evaluated and converted into raw read by the Primary Analysis Computer server. All data was automatically transferred to the Secondary Analysis Server system via the intranet. High quality sequence data was proofread and generated by PacBio’s circular consensus sequencing (CCS), then *de novo* assembled using Canu software v2.0 (Koren *et al.*, 2017), and the quality of the assembly was checked by using Quast software v5.0.2 (Gurevich *et al.*, 2013).

### Genome annotation and variant detection

SARS-CoV-2 genome sequence was annotated by VAPID v1.3 (Shean *et al.*, 2019) using the Ref_Seq Database on NCBI. Then, the sequenced was aligned with multiple sequences retrieved from GISAID and the reference sequence MN908947v3 from NCBI (Wu *et al.*, 2020) using Clustal Omega v. 1.2.4 by EMBL-EBI (Madeira *et al.*, 2019). The variants were visualized by using Geneious program v2020.1.2 (Biomatters, Auckland, New Zealand) and listed out manually.

In particular, the reference MN908947v3 is the first sequenced SARS-CoV-2 genome, originating from Wuhan, China. Other sequences were chosen in this analysis based on their location, date of sampling and divided into three groups:

1. Sequences from the Europe group:

- Initial period: Italy, February 2020.
- Pandemic period: Germany, March 2020 and England, the UK, early April 2020.
2. Sequences from the North America group:

- Initial period: Washington State (WA), USA, beginning of March 2020.
- Pandemic period: New York (NY) and Pennsylvania (PA) State, USA, late of March 2020.
3. Sequences from the Asia-Oceania group:

- Initial period: Taiwan, Vietnam, Australia and Japan (Diamond Princess cruise ship), January - February 2020.
- Pandemic period: Vietnam and Singapore, March 2020.

The Vietnamese sequences were collected from patients returing from China in January and February (Thanh Hoa, Vinh Phuc) and patients who originated from Europe, or visited the region during the spreading period in March (Hanoi, Quang Ninh).

### Variant validation by Sanger sequencing

Polymerase chain reaction (PCR) and Sanger sequencing primers were designed in-house to validate all detected variants (for more details on primer sequences used, please contact the corresponding author). PCR components for a total reaction of 25 μL were: 12.5 μL of DreamTaq Green PCR Master Mix 2X (ThermoFisher Scientific, Waltham, MA, USA), 0.5 μL of 10 pM forward and reverse primers, 100 ng of cDNA template and nuclease-free water up to 25 μL. The PCR cycling condition was set with optimized temperature for each primer set. Finally, the PCR products were separated by agarose gel electrophoresis and sequenced from both ends on the Applied Biosystems 3500 XL DNA Analyzer platform (ThermoFisher Scientific, Waltham, MA, USA) following the manufacturer’s guideline.

## RESULTS

### Isolation and cultivation of SARS-CoV-2 strain

The patient with confirmed SARS-CoV-2 was identified with cycle threshold (Ct) values of 28.06 (E gene) and 29.39 (RdRp gene). The original specimen was inoculated into cell culture. The CPE was observed 2 days post inoculation and harvested cell supernatant one day later (Figure 1). The C_t_ values of two different rRT-PCR were 15.58 for E gene and 16.68 for RdRp gene, confirming isolation of SAR-CoV-2.

**Figure 1.**
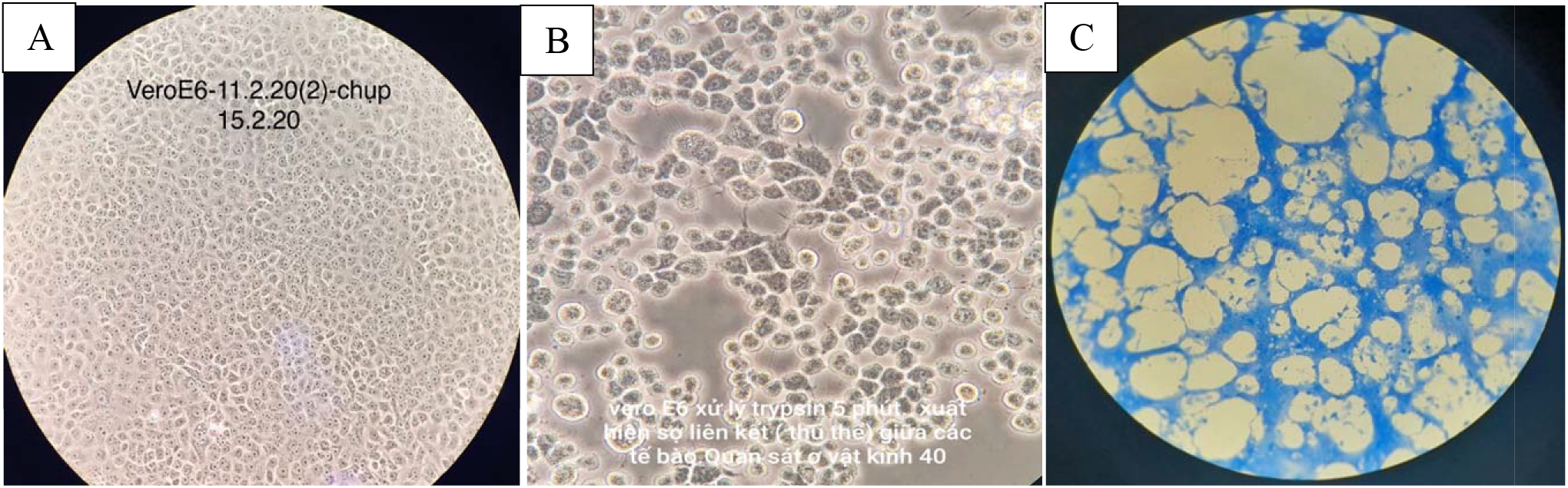
Cytopathic effects of SARS-CoV-2 in Vero E6 cell cultures. (A) Vero E6 cell cultures in negative control. (B) Vero cell lines were treated with trypsin-TPCK after 5 min. (C) CPE detachment of cells 2 days post inoculation.

### Preparation of cDNA template for whole-genome sequencing

After the evaluation of cDNA input length and quantity, we observed an average length of 1200 bp for the double stranded-cDNA, and a concentration of 1 ng/μL in total volume of 40 μL.

### Library preparation for whole-genome sequencing

The library prepared with SMRTbell Express Template Prep Kit 2.0 and Sequel Binding Kit 3.0 kit had a concentration of 1.99 ng/μL with total volume of 20 μL. Final loading concentration on the SMRT Cell was 8-pmol.

### Whole-genome sequencing on PacBio SEQUEL

The maximum reading length using chemicals and SMRT Cell version 3.0 of PacBio was approximately 140 kb; with library length of 1.2 kb, average reading length of 1.8 kb, and a N50 index of 2.1 kb. Distribution of read lengths and read quality is shown in Figure 2, showing high number of high quality (HQ) reads between 500 bp to 4000 bp (Figure 2A). Parameters that may affect the quality of the sequencing run such as adapter dimer (adapters merging instead of forming a bell with the template) and short insert (template fragments smaller than the set insert size of 1200 bp) are well within the optimal thresholds (data not shown).

**Figure 2.**
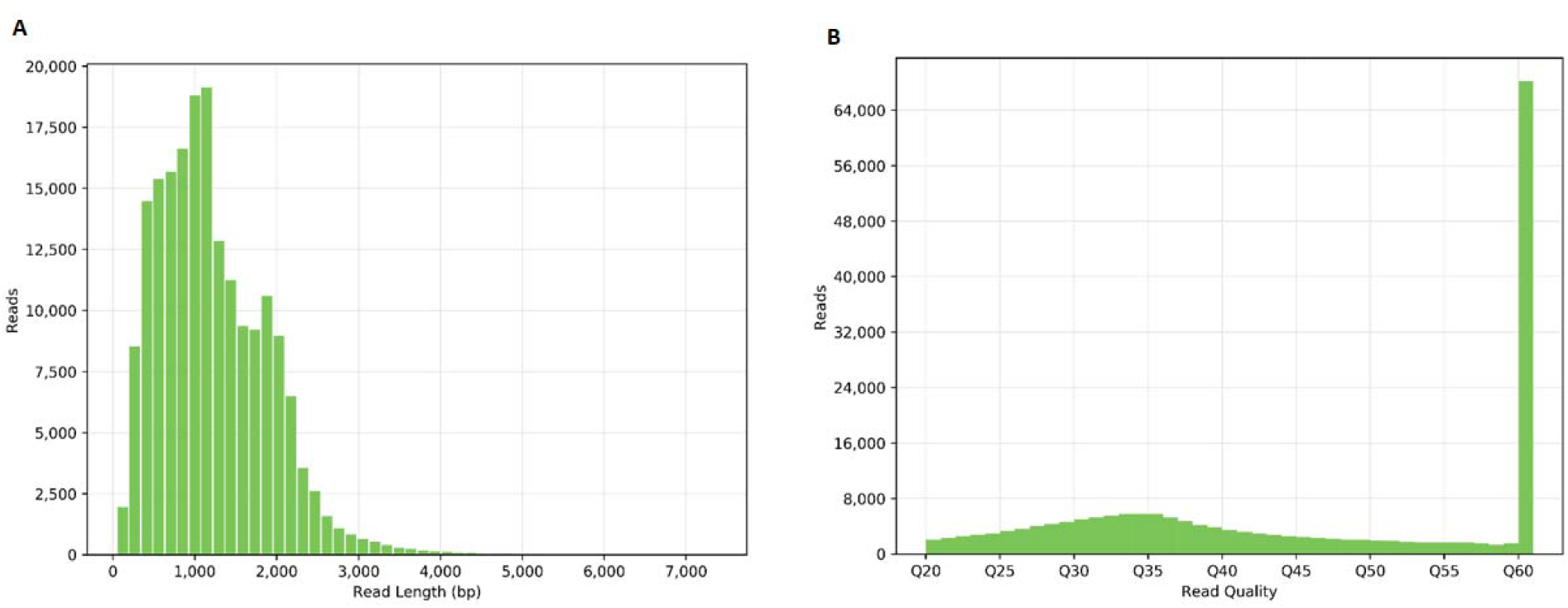
Read quality metrics of PacBio’s CCS. (A) Length distribution of high quality read. (B) Read quality distribution, from Q20 to Q60.

### *De novo* assembly and annotation of whole genome SARS-CoV-2 sequence

The total time for data quality check and processing by the Analysis Server systems was 2 hours. PacBio’s CCS results showed that the reading quality was very high: the number of >Q20 reads is greater than 192,000, resulting in more than 234 megabases (Mb) of data >Q20 with most quality reads in Q60 (Figure 2B).

Several methods for *de novo* assembly of SARS-CoV-2 whole genome sequence were tested by our group (data not shown), and Canu was the optimal software to assemble the SARS-CoV-2 genome sequence in one contig, with no gap and a length of 29,766 bp. (Table 1). VAPiD annotation results showed a complete list of 14 reported ORFs. A list of annotated ORFs as well as their locations on the sequenced viral genome in this study was shown in Table 2. The start and stop locations are different from the reference sequence MN908947; however, the length and the number of the ORFs are identical.

**Table 1.** *De novo* assembly indices of the SARS-CoV-2 genome sequence using Canu software v2.0.

| SARS_Cov_2_s.contigs | Result |
| --- | --- |
| # contigs | 1 |
| Total length | 29766 |
| Largest contig | 29766 |
| Total length | 29766 |
| GC (%) | 38.01 |
| # N's per 100 kbp | 0.00 |

**Table 3.** The annotated ORFs of SARS-CoV-2 from the *de novo* assembly sequence and their position on the virus genome.

| ORFs | Start | Stop | Length |
| --- | --- | --- | --- |
| ORF1ab polyprotein (1ab) | 247 | 21536 | 21290 |
| Surface glycoprotein (S) | 21544 | 25365 | 3822 |
| ORF3a protein (3a) | 25374 | 26201 | 828 |
| Envelope protein (E) | 26226 | 26453 | 228 |
| Membrane glycoprotein (M) | 26504 | 27172 | 669 |
| ORF6 protein (6) | 27183 | 27368 | 186 |
| ORF7 protein (7) | 27375 | 27740 | 366 |
| ORF8 protein (8) | 27875 | 28240 | 366 |
| Nucleocapsid phosphoprotein (N – 9-14) | 28225 | 29514 | 1290 |
| ORF10 protein (10) | 29539 | 29655 | 117 |

The sequence was subsequently submitted and curated on GISAID. It was given the accession ID EPI_ISL_448222 and is now available to all GISAID participants.

### Variant detection and validation

The sequence was aligned against 15 other sequences, together with MN908947v3 as the reference sequence to evaluate its accuracy and to detect any possible variant. The multiple alignment by Cluster Omega displayed 10 different single base variants compared to MN908947v3 (Table 3). The locations of these variants on the SARS-CoV-2 genome are presented in Figure 3, together with other aligned sequences.

**Figure 3.**
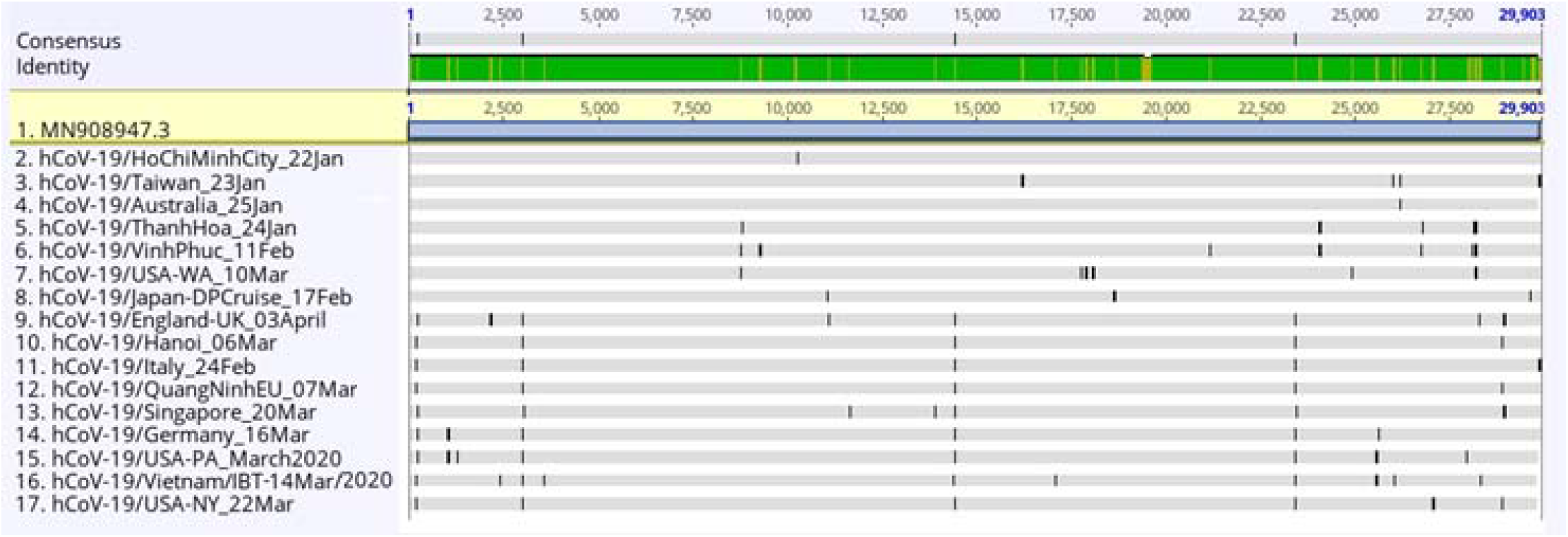
Result of the Cluster Omega multiple alignment and visualization by Geneiousv2020.1.2. Dissimilarities as compared to the reference sequence are denoted with a black strip on the gray sequence. On the identity bar, the green color indicates a 100% identity, and the copper color indicates a variant to the reference. The sequence in this study is named hCoV-19/Vietnam/IBT14-Mar/2020, at position 16. All other sequences are named with their location and date of sampling.

**Table 3.**
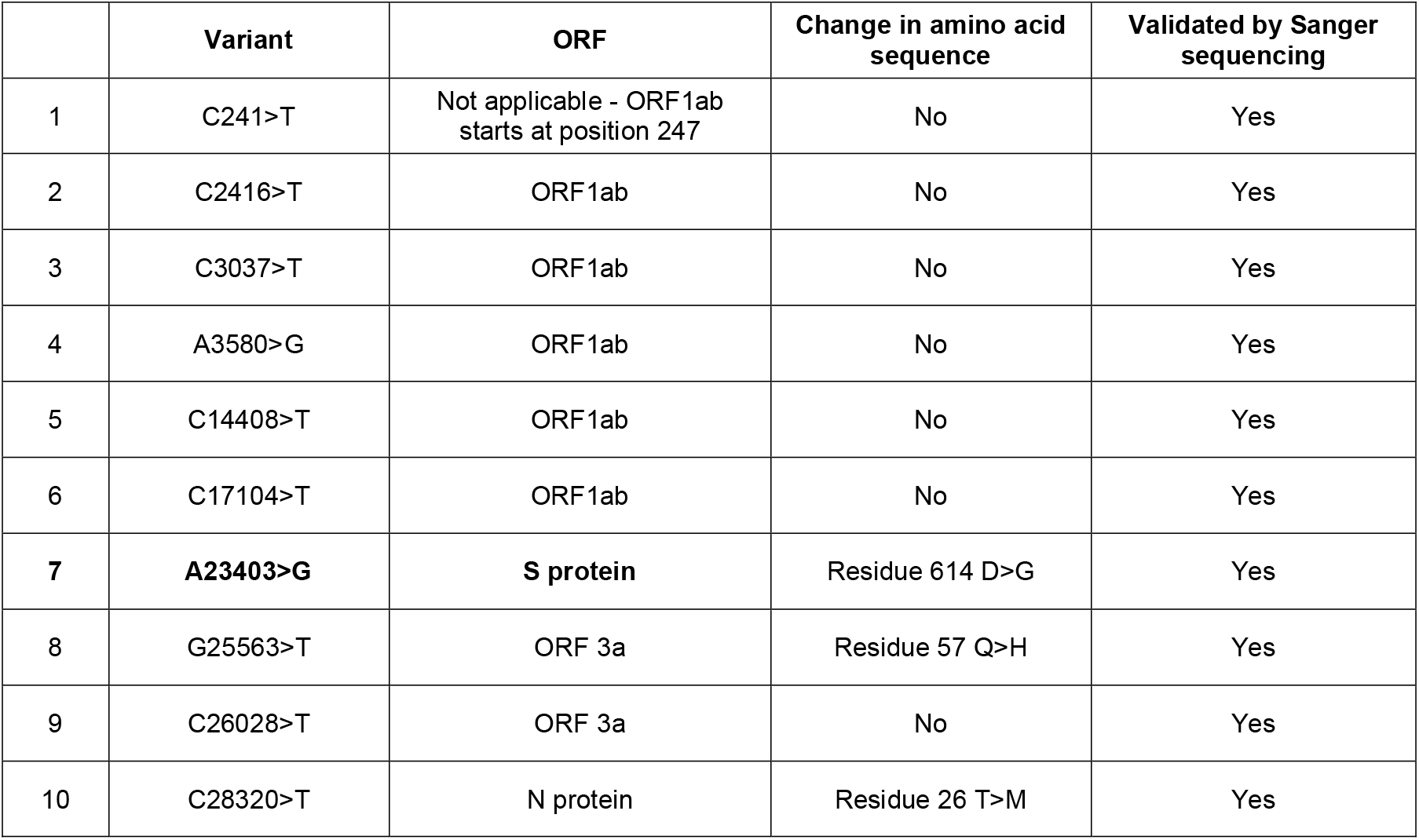
The variants detected as compared to the NCBI reference MN908947v3 using multiple alignment by Cluster Omega and visualization by Geneiousv2020.1.2.

|  | Variant | ORF | Change in amino acid sequence | Validated by Sanger sequencing |
| --- | --- | --- | --- | --- |
| 1 | C241>T | Not applicable - ORF1ab starts at position 247 | No | Yes |
| 2 | C2416>T | ORF1ab | No | Yes |
| 3 | C3037>T | ORF1ab | No | Yes |
| 4 | A3580>G | ORF1ab | No | Yes |
| 5 | C14408>T | ORF1ab | No | Yes |
| 6 | C17104>T | ORF1ab | No | Yes |
| 7 | **A23403>G** | **S protein** | Residue 614 D>G | Yes |
| 8 | G25563>T | ORF 3a | Residue 57 Q>H | Yes |
| 9 | C26028>T | ORF 3a | No | Yes |
| 10 | C28320>T | N protein | Residue 26 T>M | Yes |

Sanger sequencing has confirmed the accuracy of all variants detected in our sequence. The results of the validation by Sanger sequencing are listed in the Supplementary Data.

## DISCUSSION

The sequence in this study (sequence number 16 – Figure 3) displayed a pattern clearly shared by a group of patients who originated from Europe, or visited the region during the spreading period (sequence number 9 to 14 – Figure 3). This ‘European pattern’ includes the variants at position 241, 3037, 14408 and 23403. Among these, a notable mutation on the S protein (A23403>G, translated to a mutation from aspartic acid (D) to glycine (G) at residue 614) has been reported to be an important marker to distinguish between two viral clades in North America: the G strain is predominantly on the East Coast, and the D strain is predominantly on the West Coast (Brufsky, 2020). This mutation has also been claimed to belong to an emerging SARS-CoV-2 strain which originated from Europe (Korber *et al.*, 2020). Patient A returned from Philadelphia, Pennsylvania on the East Coast of the US. This is the region reported to be predominated by the SARS-CoV-2 G strain spread from Europe through New York (Brufsky, 2020), and our analysis has demonstrated the epidemiological evidences: the virus strain isolated from patient A and other strains from the US East Coast (Pennsylvania (15) and New York (17) – Figure 3) share the ‘European pattern’, especially the A23403>G (D614G) mutation, in their genome sequences. However, the additional variants (on ORF1ab, ORF3a and N protein) on our sequence compared to the related sequences implied an accelerated divergence of SARS-CoV-2 strains during March, when the virus was spreading aggressively globally.

In a situation where a new, unknown dangerous infectious disease can appear in the population like COVID-19, the ability to swiftly and accurately sequence the novel pathogen’s genome is critical to a country’s public health system, to provide information for research and intervention (CDC, 2020; Wellcome Sanger Insitute, 2020). Here, we have shown our capacity and preparedness for this purpose by successfully sequence and *de novo* assemble a SARS-CoV-2 whole genome in less than 30 hours, using PacBio’s SMRT sequencing technology. The genome was of high quality (Figure 2), without any gap (Table 1) and all variants called were highly accurate as confirmed by Sanger sequencing (Supplementary Data). Thus, it can fulfill many potential uses including taxonomic identification, comparative genomics, molecular test degisn for detection and development of immunological assays (Ladner *et al.*, 2014)

Notably, compared to the NCBI reference MN908947v3 with a length of 29,903 bp, this sequence is shorter by 137 bp. In particular, the 5’ terminus is 19 bp shorter, while the 3’ terminus is 118 bp shorter. However, it is important to note that the reference sequence MN908947 was sequenced using Illumina technology with a pair-end library reads of only 150 bp, resulting in a contig of 30,474 bp length from a total of 384,096 contigs assembled. Thus, it requires extra steps of RT-PCR and 5′/3′ rapid amplification of cDNA ends (RACE) to determine and confirm both the accuracy and the real length of the sequence (Wu *et al.*, 2020). Despite sharing the same problem of missing ends often encountered with *de novo* assembly (Marston *et al.*, 2013), the sequence generated by PacBio was assembled into only one contig with significantly higher coverage (1,700x compared to 600x), reducing the need to reconfirm the sequence’s accuracy.

Furthermore, the total data collected reached 11.93 gigabases (Gb) from only 8 pmol cDNA input, while the loading capacity was under the optimal threshold (data not shown), showing further potential of this system to handle dozens of SARS-CoV-2 samples on a single SMRT cell. This is crucial to any further project to mass sequence all virus strains circulating in the communities. As shown in the analysis above, SARS-CoV-2 is capable of acquiring novel mutations rapidly as it spreads in various human populations. Hence, consistent, real-time WGS data is needed to track transmission of virus, enabling the investigation of clusters and mapping how COVID-19 spreads and behaves (CDC, 2020; Wellcome Sanger Insitute, 2020). Our ability to quickly sequence highly accurate genomes combined with the potential of multiplexing dozens of samples in one run will be an asset in the efforts against not only this COVID-19 pandemic, but all future infectious disease responses.

## Acknowledgements

We would like to thank the SARS-COVID-19 response team at Institute of Biotechnology and Ho Chi Minh City Pasteur Institute. This work was financed by the Vietnam Academy of Science and Technology (VAST) Project number CT0000.05/20-20 to Prof. Truong Nam Hai as the principal investigator.

## Notes

### Competing Interest Statement

The authors have declared no competing interest.

## REFERENCE

Ashby M. Opportunities for using PacBio Long-read Sequencing for COVID-19 Research. PacBio Webinar. April 23^rd^, 2020.

Benson DA, Clark K, Karsch-Mizrachi I, Lipman DJ, Ostell J, Sayers EW (2015). GenBank. Nucleic Acids Res;43:D30–D35

Brufsky A (2020). Distinct Viral Clades of SARS-CoV-2: Implications for Modeling of Viral Spread. J Med Virol. 2020 Apr 20. doi: 10.1002/jmv.25902. [Epub ahead of print]

Centers for Disease Control and Prevention. CDC launches national viral genomics consortium to better map SARS-CoV-2 transmission. Accessed May 16th, 2020. https://www.cdc.gov/media/releases/2020/p0501-SARS-CoV-2-transmission-map.htmL

Ceraolo C, Giorgi FM (2020). Genomic variance of the 2019-nCoV coronavirus. J Med Virol. May;92(5):522–528.

Chan JF, Kok KH, Zhu Z, Chu H, To KK, Yuan S, et al. Genomic characterization of the 2019 novel human-pathogenic coronavirus isolated from a patient with atypical pneumonia after visiting Wuhan (2020). Emerg Microbes Infect.jan 28;9(1):221–236.

Chen N, Zhou M, Dong X, Qu J, Gong F, Han Y, et al (2020). Epidemiological and clinical characteristics of 99 cases of 2019 novel coronavirus pneumonia in Wuhan, China: a descriptive study. Lancet. Feb 15;395(10223):507–513.

Coronaviridae Study Group of the International Committee on Taxonomy of Viruses (2020). The species Severe acute respiratory syndrome-related coronavirus: classifying 2019-nCoV and naming it SARS-CoV-2. Nat Microbiol. Apr;5(4):536–544.

Gonzalez-Reiche Ana S., Hernandez Matthew M., Sullivan Mitchell, Ciferri Brianne, Alshammary Hala et al. (2020). Introductions and early spread of SARS-CoV-2 in the New York City area. https://doi.org/10.1101/2020.04.08.20056929 [medRxiv preprint]

Goodwin S, McPherson JD, McCombie WR (2016). Coming of age: ten years of next-generation sequencing technologies. Nat Rev Genet. May 17;17(6):333–5l.

Gurevich A, Saveliev V, Vyahhi N, Tesler G (2013). QUAST: quality assessment tool for genome assemblies. Bioinformatics. Apr 15;29(8):1072–5.

Gwinn M, MacCannell D, Armstrong GL. Next-Generation Sequencing of Infectious Pathogens. JAMA. 2019 Mar 5;321(9):893–894.

Elbe S, Buckland-Merrett G (2017). Data, disease and diplomacy: GISAID’s innovative contribution to global health. Global Challenges, 1:33–46.

Johns Hopkins University. Coronavirus COVID-19 Global Cases. Accessed May 16th, 2020. https://gisanddata.maps.arcgis.com/apps/opsdashboard/index.htmL#/bda7594740fd40299423467b48e9ecf6

Korber B, Fischer WM, Gnanakaran S, Yoon H, Theiler J, Abfalterer W, Foley B, Giorgi EE, Bhattacharya T, Parker MD, Partridge DG, Evans CM, de Silva TI, on behalf of the Sheffield COVID-19 Genomics Group, LaBranche CC, Montefiori DC (2020). Spike mutation pipeline reveals the emergence of a more transmissible form of SARS-CoV-2. doi: https://doi.org/10.1101/2020.04.29.069054 [bioRxiv preprint]

Koren S, Walenz BP, Berlin K, Miller JR, Bergman NH, Phillippy AM (2017). Canu: scalable and accurate long-read assembly via adaptive k-mer weighting and repeat separation. Genome Res. May;27(5):722–736.

Ladner JT, Beitzel B, Chain PS, Davenport MG, Donaldson EF, Frieman M, Kugelman JR, Kuhn JH, O’Rear J, Sabeti PC, Wentworth DE, Wiley MR, Yu GY; Threat Characterization Consortium, Sozhamannan S, Bradburne C, Palacios G (2014). Standards for sequencing viral genomes in the era of high-throughput sequencing. mBio. Jun 17;5(3):e01360–14.

Li Q, Guan X, Wu P, Wang X, Zhou L, Tong Y, et al. (2020). Early Transmission Dynamics in Wuhan, China, of Novel Coronavirus-Infected Pneumonia. N Engl J Med. Mar 26;382(13):1199–1207.

Lu R, Zhao X, Li J, Niu P, Yang B, Wu H, et al. (2020). Genomic characterisation and epidemiology of 2019 novel coronavirus: implications for virus origins and receptor binding. Lancet. Feb 22;395(10224):565–574.

Madeira F, Park YM, Lee J, Buso N, Gur T, Madhusoodanan N, Basutkar P, Tivey ARN, Potter SC, Finn RD, Lopez R (2019). The EMBL-EBI search and sequence analysis tools APIs in 2019. Nucleic Acids Res. Jul 2;47(W1):W636–W641.

Marston DA, McElhinney LM, Ellis RJ, Horton DL, Wise EL, Leech SL, David D, de Lamballerie X, Fooks AR (2013). Next generation sequencing of viral RNA genomes. BMC Genomics. Jul 4;14:444.

Oude Munnink BB, Münger E, Nieuwenhuijse DF, Kohl R, van der Linden A, Schapendonk CME, van der Jeugd H, Kik M, Rijks JM, Reusken CBEM, Koopmans M (2020). Genomic monitoring to understand the emergence and spread of Usutu virus in the Netherlands, 2016-2018. Sci Rep. Feb 18;10(1):2798

Quick J, Loman NJ, Duraffour S, Simpson JT, Severi E, Cowley L, Bore JA, et al. (2016). Real-time, portable genome sequencing for Ebola surveillance. Nature. Feb 11;530(7589):228–232.

Rhoads A, Au KF (2015) PacBio Sequencing and Its Applications. Genomics Proteomics Bioinformatics. 13(5):278–289.

Shendure J, Balasubramanian S, Church GM, Gilbert W, Rogers J, Schloss JA, Waterston RH (2017). DNA sequencing at 40: past, present and future. Nature. Oct 19;550(7676):345–353.

Shean RC, Makhsous N, Stoddard GD, Lin MJ, Greninger AL (2019). VAPiD: a lightweight cross-platform viral annotation pipeline and identification tool to facilitate virus genome submissions to NCBI GenBank. BMC Bioinformatics. Jan 23;20(1):48.

Vietnam Ministry of Health. Coronavirus Disease 2019 (COVID-19) Information Page. Accessed May 18th, 2020. https://ncov.moh.gov.vn/

Wellcome Sanger Insitute. UK launches whole genome sequence alliance to map spread of coronavirus. Accessed May 16th, 2020. https://www.sanger.ac.uk/news/view/uk-launches-whole-genome-sequence-alliance-map-spread-coronavirus

Wu A, Peng Y, Huang B, Ding X, Wang X, Niu P, Meng J, Zhu Z, Zhang Z, Wang J, Sheng J, Quan L, Xia Z, Tan W, Cheng G, Jiang T (2020). Genome Composition and Divergence of the Novel Coronavirus (2019-nCoV) Originating in China. Cell Host Microbe. Mar 11;27(3):325–328.

Wu F, Zhao S, Yu B, Chen YM, Wang W, Song ZG, Hu Y, Tao ZW, Tian JH, Pei YY, Yuan ML, Zhang YL, Dai FH, Liu Y, Wang QM, Zheng JJ, Xu L, Holmes EC, Zhang YZ (2020). A new coronavirus associated with human respiratory disease in China. Nature. Mar;579(7798):265–269.

Zhou P, Yang XL, Wang XG, Hu B, Zhang L, Zhang W, Si HR, Zhu Y, Li B, Huang CL, Chen HD, Chen J, Luo Y, Guo H, Jiang RD, Liu MQ, Chen Y, Shen XR, Wang X, Zheng XS, Zhao K, Chen QJ, Deng F, Liu LL, Yan B, Zhan FX, Wang YY, Xiao GF, Shi ZL. A pneumonia outbreak associated with a new coronavirus of probable bat origin. Nature. 2020 Mar;579(7798):270–273.

