## Supplementary figures and images for "WHOLE-GENOME SEQUENCING AND *DE NOVO* ASSEMBLY OF A 2019 NOVEL CORONAVIRUS (SARS-COV-2) STRAIN ISOLATED IN VIETNAM"

### Supplementary Data

**Supplementary Data**

**Results of variant-calling validation by Sanger sequencing.**

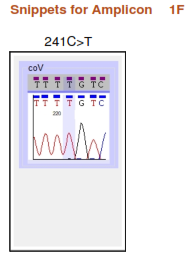

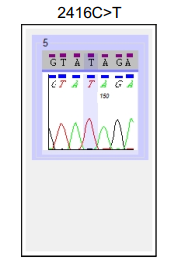

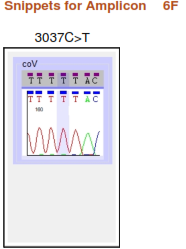

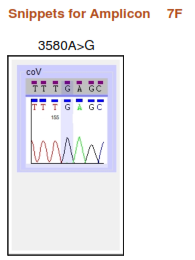

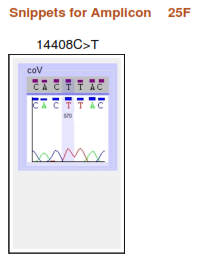

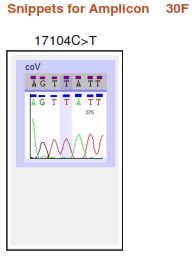

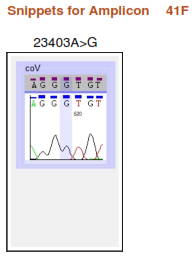

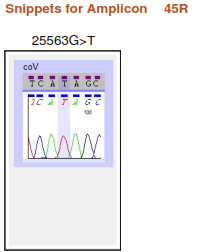

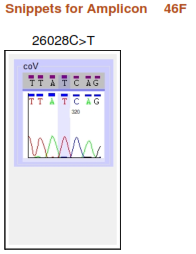

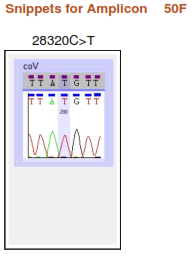
